# How Does Transcription-Associated Mutagenesis Shape tRNA Microevolution?

**DOI:** 10.1101/2025.03.26.645593

**Authors:** Hector Baños, Ling Wang, Corinne Simonti, Annalise Paaby, Christine Heitsch

## Abstract

Transfer RNAs (tRNAs) are among the most highly conserved and frequently transcribed genes. Recent studies have demonstrated that tRNAs experience exceptionally high rates of transcription-associated mutagenesis (TAM) as well as strong purifying selection. How the mutational input of TAM, which induces a non-uniform distribution of nucleotide substitutions, affects the fitness of tRNA molecules is unclear. Secondary structure in tRNAs is strongly conserved over macro-evolutionary time, suggesting that mutations that disrupt paired sites may be especially deleterious, but TAM-induced mutations primarily involve nucleotide transitions, which tend to preserve base-pairing stability.

To examine how TAM affects tRNA molecule fitness and shapes tRNA evolution over short timescales, we analyzed tRNA allelic variation in contemporary *Caenorhabditis elegans* strains. We propose a model of tRNA microevolution driven by TAM and demonstrate that the observed secondary structure characteristics align with our predicted TAM-biased patterns. Furthermore, we developed a continuous Markov substitution model that incorporates TAM-specific mutational biases. This TAM-biased model fits the *C. elegans* tRNA data more effectively than standard models, such as the general time-reversible (GTR) model.

Based on these results, we conclude that tRNAs in natural populations carry substantial levels of structure-destabilizing mutations, which may be tolerated but nevertheless likely induce meaningful fitness costs. Our findings are consistent with recent experimental studies on tRNA fitness in yeast but challenge prior theoretical and computational analyses that emphasize RNA base-pairing as a primary determinant in genotype-phenotype systems.

**Significance Statement:** Transfer RNAs (tRNAs) are ancient molecules, encoded as genes in all living systems. tRNA genes are known to experience exceptionally high rates of both mutation and purifying selection, but how these opposing evolutionary forces shape tRNA evolution is unclear. We developed a sequence substitution model specific to tRNA mutagenesis and applied it to standing variation in a natural population in order to infer how mutation and selection affect the structural stability of tRNA molecules.

## 2 Introduction

Transfer RNAs (tRNAs) are ancient and indispensable genes essential for protein synthesis, renowned for their evolutionary conservation (Tang et al., 2009; Zhang and Ferré-D’Amaré, 2016). This conservation is widely attributed to the structural integrity necessary for their function (Westhof et al., 2022), maintained by strong selection for base-pair preservation. As a result, the preservation of secondary structure is frequently used as a proxy for tRNA functionality, making tRNAs a classic model for exploring genotype-phenotype relationships.

However, the evolutionary forces shaping tRNA sequences remain poorly understood, particularly at the population level (Ishimura et al., 2014). The preservation of secondary structure over macroevolutionary time emphasizes its importance to molecular function, but tRNA sequence confers essential functionality beyond the context of folding. For example, sequence identity elements specify interactions with aminoacyl-tRNA synthetases and other factors necessary for appropriate amino acid charging (Giegé and Eriani, 2023). Recent yeast experiments show that single mutations in loop regions can significantly affect fitness, while single mutations in the acceptor stem are surprisingly well-tolerated (Li et al., 2016). These findings provide the first empirical estimates of tRNA fitness effects and challenge the prioritization of paired sites over loop regions implied by the thermodynamic model (Fontana and Schuster, 1998; Reidys et al., 2001; Aguirre et al., 2011). Alterations to sequence identity elements may disrupt co-adaptations that arise over relatively short timescales (Meiklejohn et al., 2013; Adrion et al., 2015), but the fitness dynamics of tRNAs over microevolutionary time, including how selection shapes the distribution of standing variants in populations, remains unknown.

Transcription-associated mutagenesis (TAM) (Jinks-Robertson and Bhagwat, 2014) has been identified as a major contributor to mutation rates in highly transcribed regions, and tRNAs are among the most highly transcribed genes in the genome (Palazzo and Lee, 2015; Boivin et al., 2018). During transcription, deamination of the noncoding strand, along with DNA repair mechanisms responding to this deamination, is associated with an increase in *C* → *T* and, secondarily, *G* → *A* mutations on the coding strand (Green et al., 2003; Jinks-Robertson and Bhagwat, 2014). tRNAs within and across species exhibit mutational variation with signatures of historical TAM, including in flanking regions just upstream and downstream of tRNA gene bodies (Thornlow et al., 2018). Both the gene and its flanking regions are vulnerable to TAM during transcription, and are presumed to experience the same rate of mutation. However, strong purifying selection on the gene itself purges some mutations, while flanking regions can accumulate exceedingly high numbers of mutations. Thus, these regions provide a record of the historical expression level and mutation rate of that gene (Thornlow et al., 2018, 2020).

The exceedingly high rates of mutation and purifying selection at tRNAs has led to the hypothesis that eukaryotic genomes carry substantial mutational load at tRNA loci (Thornlow et al., 2018). However, how this translates to a disease burden, including how the fitness of individual tRNA molecules are compromised, remains unexplored. A key question is how the non-uniform nucleotide substitutions associated with TAM might alleviate or exacerbate destabilization of tRNA secondary structure. For example, selection might tolerate transitions over transversions, if they are less likely to disrupt nucleotide pairs in stems; under this scenario, the *C* → *T* and *G* → *A* transitions induced most frequently by TAM may not be as deleterious as they could be. Also under this scenario, the coincidence of the highest frequency and least deleterious mutations would mean that selection would not erode the TAM signature from standing variation even as most mutations are purged (Thornlow et al., 2018). Our focus is to better assess the signature of TAM in tRNA allelic variation and determine how TAM affects overall tRNA fitness.

It is important to note that beyond TAM, additional mechanisms govern the occurrence of mutations that comprise standing variation in tRNA genes, including transcription-coupled repair (Svejstrup, 2002). However, a strong and growing body of literature indicates that highly expressed genes are subject to higher rates of mutation, and that TAM dominates the mutational landscape of tRNA genes (Park et al., 2012; Jinks-Robertson and Bhagwat, 2014; Liu and Zhang, 2020; Thornlow et al., 2018, 2020).

In this study, we assess how TAM affects tRNA fitness and microevolution by analyzing standing genetic variation in tRNA genes in *Caenorhabditis elegans*. Using a novel continuous Markov substitution model, we evaluate TAM’s mutational signature and the role of selection on base-pair preservation. Aligned with the experimental results in (Li et al., 2016; Domingo et al., 2018), the data, supported by our model, shows that paired sites experience higher substitution frequencies than unpaired ones under TAM. Additionally, we observed that most substitutions, whether preserving base-pairing or not, decrease thermodynamic stability, suggesting that TAM compromises structural integrity rather than maintaining it. These results highlight how TAM shapes tRNA evolution in ways beyond structural constraints under the thermodynamic model.

## 3 Background

Markov models are widely used to study evolutionary processes, particularly DNA substitutions (Felsenstein, 2003; Yang, 2006). While these models are traditionally applied to gene tree inference across species, our approach departs from this by using them to describe allele formation within a population.

These models use different parameters to capture the various mutational mechanisms and selection pressures shaping DNA substitutions. Hence, the Markov exchangeability matrix for the substitution from *i* to *j* (denoted *i* → *j*) for *i, j* ∈ {*A, C, G, T* } may differ depending on the combination of nucleotides. In particular, transitions are expected to occur much more frequently than transversions since they preserve the purine (*A* ↔ *G*) versus pyrimidine (*C* ↔ *T*) distinction.

Moreover, for noncoding RNA genes, it is likely that transitions are further enhanced as they allow Watson-Crick base pairs G–C or C–G (denoted GC) and A–U/U–A (AU)^a^ to inter-convert via the wobble G–U/U–G (GU) pairing:

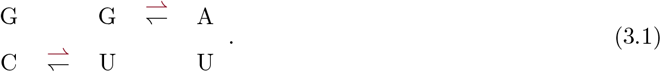

Note that of the three types of canonical base pairings, GC is the most thermodynamically favorable, followed by AU, and then GU. Hence, based on the pairing conservation, one would expect *C* → *T* transitions to be favored in GC pairs, *A* → *G* ones in AU, and both *G* → *A* and *T* → *C* in GU with the last as the most preferred, and the first as the least, due to thermodynamic stability. TAM is the dominant source of mutations at tRNAs (Thornlow et al., 2018), which is biased towards more frequent *C* → *T* and *G* → *A* transitions on the coding strand, denoted in red above.

### 3.1 Secondary structure conservation in RNA macroevolution

A noncoding RNA molecule like tRNA is composed of a single-stranded sequence which folds hierarchically (Tinoco and Bustamante, 1999) into a functional structure. The intra-sequence base pairings of the secondary structure are the critical scaffold which is then organized by tertiary interactions into the final 3D conformation. In some cases, it is the pattern of base pairing, and not necessarily the nucleotide content, that is evolutionarily conserved. This selection pressure can be so strong that the occurrence of compensatory mutations in base pairs, identified under multiple sequence alignment, is the gold-standard for RNA structural inference (Eddy and Durbin, 1994; Cannone et al., 2002; Griffiths-Jones et al., 2005).

More precisely, a *secondary structure* for an RNA sequence *R* of length *n* is a set of paired indices *S*(*R*) = {(*i, j*) | 1 ≤ *i < j* ≤ *n*} such that the nucleotides in positions *i* and *j* form a canonical base pair, i.e. either Watson-Crick (GC, AU) or wobble (GU). Suppose that *R*^*′*^ is a *one-point mutant* of *R*, meaning that it differs from *R* by a single substitution. Suppose further that *S*(*R*) is also a valid secondary structure for *R*^*′*^ (meaning that the nucleotide mutation in *R*^*′*^ does not disrupt any pairing of *S*(*R*)). Then either the substitution changed an unpaired nucleotide or it changed a paired one as described in Equation (3.1). In these cases we say that the substitution *preserves pairing potential*. Otherwise, it necessarily disrupts the pairing involving the mutated nucleotide.

It is critical to emphasize that a substitution being pairing potential preserving says nothing about the secondary structure of *R*^*′*^. The mutational constraint only requires that *R*^*′*^ *can* assume the same set of pairings as *S*(*R*), not that it *does*. It is, however, a necessary condition for *R*^*′*^ to be a *neutral neighbor* of *R*.

Using RNA secondary structure formation as a model genotype-phenotype system (Schuster et al., 1994; Cowperthwaite and Meyers, 2007), the theory of neutral networks was developed as a framework for understanding structural fitness landscapes. In this context, evolution is modeled as a series of one-point mutations, and a substitution is considered “neutral” if it does not alter the secondary structure. The conservation of pairing (and not just its potential preservation) is most often determined by free energy minimization (Zuker and Stiegler, 1981) under the nearest neighbor thermodyanmic model (NNTM). Hence *R*^*′*^ is a neutral neighbor of *R* if it differs by a single substitution and retains the same minimum free energy (MFE) structure, i.e. the one-point mutation must be *MFE-preserving*.

Under this NNTM optimization interpretation of fitness neutrality, disruption of canonical base pairs is highly deleterious. Hence the macroevolution expectation of secondary structure conservation implies that substitutions should follow these patterns:

(U) A bias against mutations at paired sites.

(P1) A bias towards *C* → *T* in GC pairs and *A* → *G* in AU ones if a Watson-Crick pairing is mutated.

(P2) A bias towards transitions if a wobble pairing is mutated, with *T* → *C* clearly favored over *G* → *A*.

### 3.2 Experimental insights into tRNA microevolution

Transfer RNA has one of the most direct genotype-phenotype relationships possible due to the tight coupling of sequence/structure/function. In order to read the mRNA codon and deliver the amino acid, a tRNA sequence must fold into a canonical three-dimensional ‘L’ shape. This essential structure is remarkably conserved across all three domains of life (Daewoo Pak and Burton, 2017). Although variations exist– including mitochondrial tRNAs that adopt extreme, yet functional shapes (Jühling et al., 2018; Ozerova et al., 2024), the cloverleaf secondary structure remains the core scaffold of this critical arrangement. As illustrated^b^ in Figure 1, the intra-sequence base pairing forms four runs of stacked base pairs, with three hairpin loops and the central multibranch loop. Three nucleotides in the anticodon loop target the complementary codon in the mRNA molecule while the corresponding amino acid, attached at the Tail, is loaded on the opposing acceptor stem. As pictured, the variable portion (M3) of the multiloop is unpaired, but it can sometimes be much longer (Berg and Brandl, 2021) in which case it typically includes pairings.

**Figure 1.**
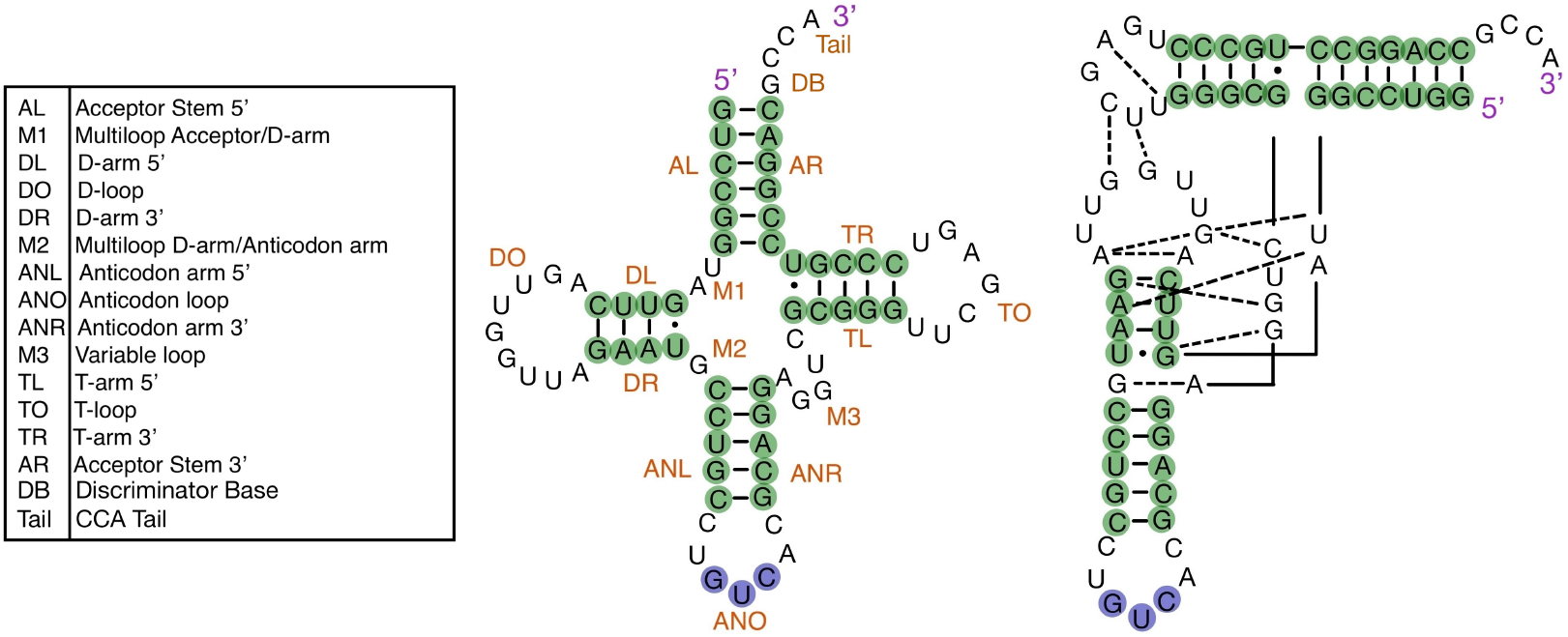
Representative tRNA secondary structure and tertiary interactions. From left to right: Table lists the structural components, including the CCA Tail, which are then labeled on the secondary structure. The four-armed cloverleaf is closed by the acceptor (A) stem, and contains three hairpin stem-loop structures: the D, anticodon (AN), and T arms. Paired nucleotides (green) as well as the anticodon (blue) are highlighted. Most pairings are Watson-Crick (GC/AU, dash) but two wobble ones (GU, dot) are present. Various tertiary interactions (dashed lines) stabilize the 3D structure. The overall L-shape is critical to ribosome binding, and hence protein biosynthesis.

As a model genotype-phenotype system, tRNA folding has attracted considerable theoretical attention (Fontana and Schuster, 1998; Reidys et al., 2001; Aguirre et al., 2011) in macroevolution studies. These studies use secondary structure thermodyanmic predictions, such as the *RNAfold* function of the ViennaRNA 2.0 package (Lorenz et al., 2011) which outputs a single MFE structure, as a proxy for evolutionary fitness. In this neutral-neighbor context, any change in the base pairing pattern, i.e. the positions which are paired, is deemed deleterious.

More recently, however, tRNA has been the focus of growth experiments (Yona et al., 2013; Li et al., 2016; Domingo et al., 2018) that use sequencing technology to explore fitness at the microevolution scale. All three studies investigated a particular *Saccharomyces cerevisiae* Arginine gene: 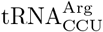, which is distinctive^c^ in having a single copy with anticodon CCU. Hence, because there is only one instance in the genome, changes in growth rate can be directly attributed to changes in this gene.

By deleting the gene entirely, Yona et al. (2013) demonstrated that within 200 generations the wild-type growth rate could be recovered. Investigating further they confirmed that this was achieved by mutating one of the 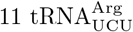 copies available, and not always the same one. Hence, a single *C* → *T* substitution in the wobble position of the anticodon was sufficient to recover the wild-type fitness. Moreover, they then confirmed that such anticodon switching can be found across all three domains of life, leading to the conclusion that this is a wide-spread adaptation mechanism. Hence, while the anticodon is generally expected to be under very strong selection pressure, it can and will evolve rapidly to meet new translation demands.

In contrast, Li et al. (2016) comprehensively characterized all one-point mutants of 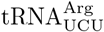. They classified 1% as beneficial, 42% as deleterious, and the remaining majority (57%) as “(nearly) neutral.” Generally, the least mutable positions were in loops, and the most in stems. As expected, mutations in the anticodon itself significantly reduced fitness. Nonetheless, mutations at three positions in the T-loop (positions 53, 54, and 55), two in the T-stem (positions 52 and 60, which form a the base pair ‘closing’ the T-loop), and one in the D-stem also had a substantial negative impact (position 18). Conversely, the acceptor stem seemed to tolerate all single substitutions, as did the variable loop.

This robustness to one-point mutants in key structural components was echoed by Domingo et al. (2018) who investigated all existing variants of 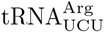 found across yeast species. They identified 14 mutations across 10 sites in 6 other genomes: 11 in the acceptor stem (5 in AL, 6 in AR) and 2 in the variable loop (M3) with 1 from the anticodon stem (ANL). All but one of the AL/AR mutations came together in compensatory pairs, of which 1 was a GU, 3 AU, and 1 GC. The remaining AR mutation formed a GU pair as did the ANL one. They then generated all possible combinations of these substitutions and tested their fitness in growth competition. All 14 one-point mutants exhibited fitness comparable to the wild-type *S. cerevisiae* gene and the extant variants.

Both studies found that fitness decreased rapidly with subsequent mutations, with significant epistatic interactions also identified. The epistasis had a strong negative bias except — unsurprisingly — for compensatory mutations in paired positions. All of this suggests that, while conservation of secondary structure is critical at the macroevolutionary level, the situation may be much more nuanced and flexible across a population.

### 3.3 Markov models for DNA macroevolution

DNA sequences evolving in time under nucleotide substitutions are typically modeled using a continuous-time Markov process. These models are parameterized by a nucleotide distribution vector *π* = (*π*_*A*_, *π*_*C*_, *π*_*G*_, *π*_*T*_) and 4 *×* 4 instantaneous rate matrix *Q*. The Markov matrix *M* of the model is defined as *M* = exp(*Qt*), where the entry *m*_*ij*_ ∈ *M* denotes the probability that nucleotide *i* mutates to nucleotide *j* after time *t*. Then the joint distribution of nucleotides in the ancestral and descendant sequences over time *t* is given by *P* = diag(*π*) exp(*Qt*) where the entry *p*_*ij*_ represents the probability of observing a substitution *i* → *j* at a site.

In general, the models are distinguished by the constraints on *π* and *Q*. All forms require that ∑*π*_*i*_ = 1 with *π*_*i*_ ≥ 0, that *q*_*ij*_ *>* 0 for *i* ≠ *j*, and that *q*_*ii*_ = − ∑ _*i*_ ≠ _*j*_ *q*_*ij*_. In other words, *π* must be a probability distribution, all off-diagonal entries of *Q* are positive, and the diagonal ones make the row sums equal zero. In a context where the edge length is unknown, one off diagonal entry is set to 1, to avoid scaling redundancy. Additionally, the rate matrix used is typically normalized by multiplying all original entries of *Q* by

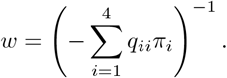

This results in the expected number of substitutions being one per unit of time.

The earliest, and most mathematically tractable models, assume that *π* is uniform, *t* is unknown, and the 11 free entries of *Q* come from a set of either 0, 1, or 2 parameters (Jukes and Cantor, 1969; Kimura, 1980). When there are two, one is for transitions (which preserve the purine/pyrimidine division) and the other for transversions. The addition of a third parameter is used to distinguish transversions which are amino/keto preserving from the weak/strong symmetry.

More biologically realistic models allow nonuniform *π* and provide a larger number of exchangeability parameters. The most general form, associated with the *continuous general Markov model* (GM) over a single edge, has no additional constraints, and hence 12 free parameters (11 from the rate matrix, and 1 from the edge length). However, this great flexibility often results in over-fitting, and is seldom needed to capture the substitution process accurately.

The most common substitution model used for phylogenetic inference is the *general time-reversible* (*GTR*) one (Tavaré, 1986). Such model has 5 free parameters (6 when considering a single edge length) satisfying diag(*π*)*Q* = *Q*^*T*^ diag(*π*). This equality holds when the substitution process has the same parameters going forward in time as backward. In a GTR model, the rate matrix *Q* can be parameterized via a 4 *×* 4 non-negative matrix known as and exchangeability matrix. As useful as the assumption of time-reversibility is for macroevolution analyses, it is not clear that it will be appropriate for the *C. elegans* tRNA population data. Although we considered *GTR* as a possibility, our TAM-biased Markov substitution model is built from the strand-symmetric rate matrix. We note that a model can be both strand-symmetric and time-reversible only if *π*_*A*_ = *π*_*T*_ and *π*_*C*_ = *π*_*G*_.

The strand-symmetric (*sym*) model (Casanellas and Sullivant, 2005) also has 6 free parameters. However, the equality now enforced is on the exchangeability of *i* → *j* and its Watson-Crick complement, i.e. *C* → *T* and *G* → *A* must have the same parameter. The premise is that, for many mutational mechanisms, it is not possible to determine whether a substitution initially occurred in the coding strand or the template one. We consider such mutations as background noise, and take this matrix as the foundational one for our new TAM-biased model.

### 3.4 Microevolution signature of mutational bias

Transfer RNAs are not only highly conserved but also highly transcribed, exposing them to high levels of *transcription-associated mutagenesis* (TAM) (Thornlow et al., 2018). Critically, TAM’s characteristic signature breaks the strand-symmetry assumption, as well as the time-reversibility one.

During RNA transcription, a hybrid DNA–RNA complex forms, leaving the non-template DNA strand exposed and vulnerable to mutagens (Jinks-Robertson and Bhagwat, 2014). The formation of non-canonical structures by the unwound DNA further promotes mutagenesis by inhibiting repair pathways and interfering with replication processes (Gaillard and Aguilera, 2016). Thus, the more frequently a gene is transcribed, the more vulnerable its coding strand becomes to mutations. TAM induces a non-uniform, biased distribution of mutations, leading to a high frequency of *C* → *T* and, secondarily, *G* → *A* mutations, on the coding strand (Thornlow et al., 2018; Jinks-Robertson and Bhagwat, 2014). For example, and mentioned in Gómez-González and Aguilera (2007); Jinks-Robertson and Bhagwat (2014); Thornlow et al. (2018), activation-induced cytidine deaminase (a notable source of TAM) mutates DNA during transcription by changing cytosines to uracils, resulting in *C* → *T* and *G* → *A* substitutions. To fix the base-pair mismatch, the opposing *G* is converted to *A* and *U* to *T*, resulting in excess *C* → *T* mutations on the non-template strand and excess *G* → *A* mutations on the template strand. Since the non-transcribed strand is more exposed as single-stranded DNA within the transcription bubble, such strand is more susceptible to cytosine deamination, leading to more *C* → *T* relative to *G* → *A* substitutions (Williams et al., 2023).

The discovery that tRNAs are subject to exceptionally high levels of TAM also demonstrated that tRNAs experience very strong purifying selection (Thornlow et al., 2018). This is evident by the reduced level of mutational variation within tRNA gene bodies relative to their flanking regions. Both tRNAs and their flanking regions presumably acquire a similar spectrum of mutations, but those deleterious changes to tRNA that are purged are not observed in population data. In this way, the mutational variation at tRNA flanking regions offers a relatively direct record of the historical TAM process on the gene itself. Here, we use this unique phenomenon to estimate the mutational input to tRNAs from flanking sequence data, and test hypotheses about how TAM affects tRNA fitness and how tRNAs respond to selection.

## 4 Results and Discussion

### 4.1 Model validation

We find good agreement between model hypotheses and population data. Specifically, the new mutational variation sequence *S*^*′*^ reasonably describes the observed tRNA distribution, and the consensus sequence *S* is a suitable approximation to the most recent common ancestor of those *C. elegans* alleles. Moreover, the flanking regions yield a robust estimate for the TAM bias vector *τ*, and the tRNA distribution of substituted nucleotides in the tRNA (that is, the proportion of nucleotides that mutated to another nucleotide) align with *π*^*τ*^, the proposed TAM biased distribution. Finally, of the Markov substitution models considered, the new *TAM* matrix best describes the data.

Our full dataset consisted of 661 variants for 581 tRNA genes. Of those alleles, 70 have an indel and cannot be described by a substitution model. Among the 591 remaining, no two differ at the same position, but 60 have two or more substitutions. Hence, there are 531 distinct-site, one-point mutant alleles, and the compound sequence *S*^*′*^ encodes 80.3% of the total tRNA mutational variation in the *C. elegans* population.

We consider the evolution of *S*^*′*^ from *S*, the consensus sequence for the 531 genes. This is a good approximation to their most recent common ancestor *S* because the distribution of alleles is very sparse. In particular, nearly all consensus tRNA (484*/*531 = 91.1%) occur in nearly all strains (at least 298*/*331 so ≥ 90.0%). The remaining 47 consensus sequences have more frequently occurring variants, including missing entirely. However, of the 45 present in ≥ 45% of strains, only 2 have a variant which occurs nearly as often (*<* 15% difference). Hence, in all but 4 tRNA genes, the consensus clearly dominates.

The TAM bias vector *τ* was estimated from the tRNA flanking regions, which contained 2070 mutations. Flanking sequence are subject to TAM but not to direct selection, so they provide a putatively unfiltered empirical estimate of the distribution of input mutations. This yielded *π*−*π*^*τ*^ = (0.0454, −0.1111, −0.0456, 0.1110) = (*τ*_*R*_, −*τ*_*Y*_, −*τ*_*R*_, *τ*_*Y*_). The close match in positive and negative estimates supports our TAM bias hypothesis; averaging yields 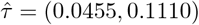.

The nucleotides A/C/G/T in *S* are distributed as *π* = (0.181, 0.258, 0.323, 0.237)^d^. Under our TAM bias model, the expected distribution of substituted nucleotides is *π*^*τ*^ = (0.135, 0.370, 0.369, 0.125) while 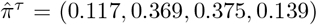 is observed. Under statistical testing, described in *Materials and Methods*, we fail to reject the null hypothesis with a *p*-value of 0.510 whereas repeating the analysis with *τ* = (0, 0) yields *p <<* 0.001. Hence we conclude that the tRNA substitutions in the data are well described by our proposed TAM bias distribution.

This distribution of substituted nucleotides is then used to parameterize a new rate matrix, denoted *TAM*, for the continuous Markov substitution process model. As illustrated in Figure 2, *TAM* is a special case of *tran*, which was introduced as a generalization of the standard *sym*. We also consider the full *GM* as well as the frequently-used *GTR*.

**Figure 2.**
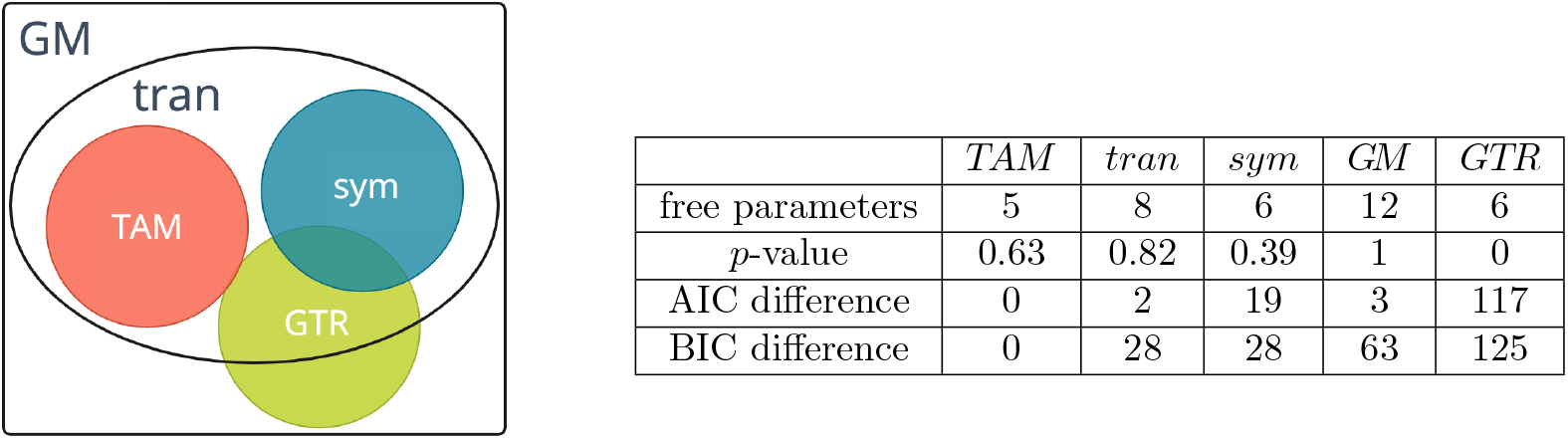
Comparison of rate matrix parameterizations for Markov substitution model: general Markov (*GM*), strand-symmetric (*sym*), transition-asymmetric (*tran*), and TAM-biased (*TAM*) along with general time-reversible (*GTR*). The Akaike and Bayesian information criteria (AIC, BIC) are reported as the difference from *TAM* ‘s score as this was the lowest, i.e. best.

The calculated *p*-values, along with other model selection criteria, are also given in Figure 2. With *p <<* 0.001, it is clear that *GTR* does not fit the data. Having rejected the null hypothesis, and in view of the significantly lower AIC and BIC scores, we conclude that time-reversibility — a common macroevolution assumption — is not an appropriate expectation for this microevolution model.

Among the remaining four matrices with *p >* 0.05, the model selection criteria support *TAM* as the best fit for the substitution process which model the formation of *S*^*′*^ from *S. TAM* has the lowest AIC, although the difference with *tran* is on the borderline of meaningful. (Recall that differences of less than 2 units in AIC and 6 in BIC are generally considered negligible.) However, BIC is definitive; *TAM* had the lowest (best) score by a wide margin. Moreover, there is no reason to prefer *tran* over *TAM* as the latter provides more specific biological insights about the distribution of substituted nucleotides that the former does not.

Finally, we note that the *GM p*-value likely indicates overfitting, and yet the information criteria support *TAM* as a better fit to the *C. elegans* tRNA data.

### 4.2 Assessing preservation of pairing potential

As part of the model validation, we confirmed that the observed substitutions in the population follow a distribution consistent with the TAM bias expected by the model. In particular, C’s are overrepresented in 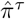 by 43.0% while G’s increased by 16.0% over the initial nucleotide distribution *π*. This demonstrates that the mutational variation in extant tRNAs exhibits a signature consistent with TAM, even though these sequences have also experienced strong selection that has removed many mutations over time.

We now assess whether three related distributions — nucleotides by pairedness (Π), which separates *π* into paired and unpaired sites; pairedness (Π_*pu*_), the pairedness distribution achieved by column marginalization of Π; and base pairs (*ρ*), which indicates the proportion of GC, AU and GU pairings in the secondary structure — also follow TAM-biased distributions. Details are give in Table 1. In all three cases, we fail to reject the null hypothesis for the expected and observed distributions of substituted nucleotides (i.e., paired sites also show signature of TAM).

**Table 1.**
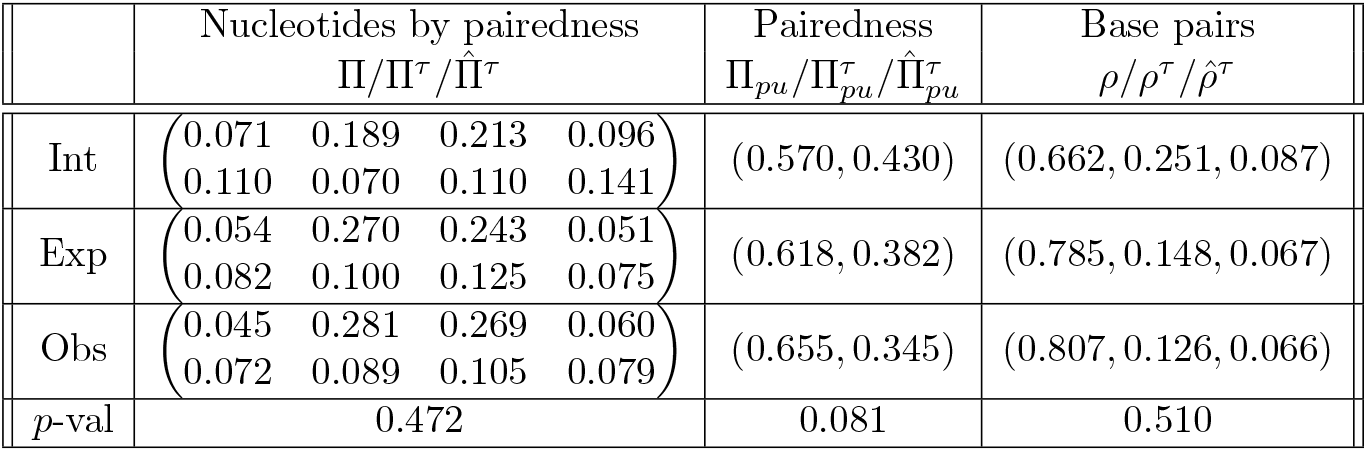
Comparison of three expected pairing distributions under TAM bias model given initial ones from *S* and estimated 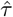 with observed alleles encoded in *S*^*′*^.

In fact, as seen in the first and third columns, the proportion of C and G nucleotides in base pairs which are mutated is consistently *higher* than expected. That is, even though the TAM bias leads us to expect more C and G substitutions, we observed even more than the model predicted. While this is not statistically significant, the trend, if representative, is surprising as it is inconsistent with expectations for the preservation of base pair formation.

One expectation for pairing preservation is that unpaired nucleotides should better tolerate mutations, and therefore exhibit more substitutions in the dataset. This is clearly violated as paired positions are mutated almost twice as often (0.655 versus 0.345). Specifically, this represents an increase of 14.9% over the initial Π_*pu*_ one. As seen when 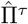 is compared to Π, this is entirely due to more substitutions in paired C’s and G’s which increased by 48.8% and 26.2% respectively over the initial distribution. This is consistent with a TAM-biased substitution process. That is, paired nucleotides contain more C and G nucleotides, which makes these sites more likely to be targeted by TAM.

From the selective pressure to preserve secondary structure, one might then expect that mutated GC pairs are predominantly converted to GU pairings under the *C* → *T* transition as these would preserve structure in a wobble pair (Thornlow et al., 2018). As seen in Figure 3, this is indeed the most common substitution. However, it happens only 33.5% of the time compared to the destabilizing *G* → *A* transition and the four relevant transversions. Likewise, an AU pairing is potentially preserved as a GU one in only 31.8% of substitutions. However, these proportions are actually lower than when the corresponding unpaired substituted nucleotides combinations are considered: 47.1% for C,G and 45.3% for A,U. Hence, in this microevolution setting, we do not see strong support for selection favoring the formation of GU pairs from either GC or AU ones. Therefore, we see that *C* → *T* mutations are not preferentially biased to maintain structural conservation.

**Figure 3.**
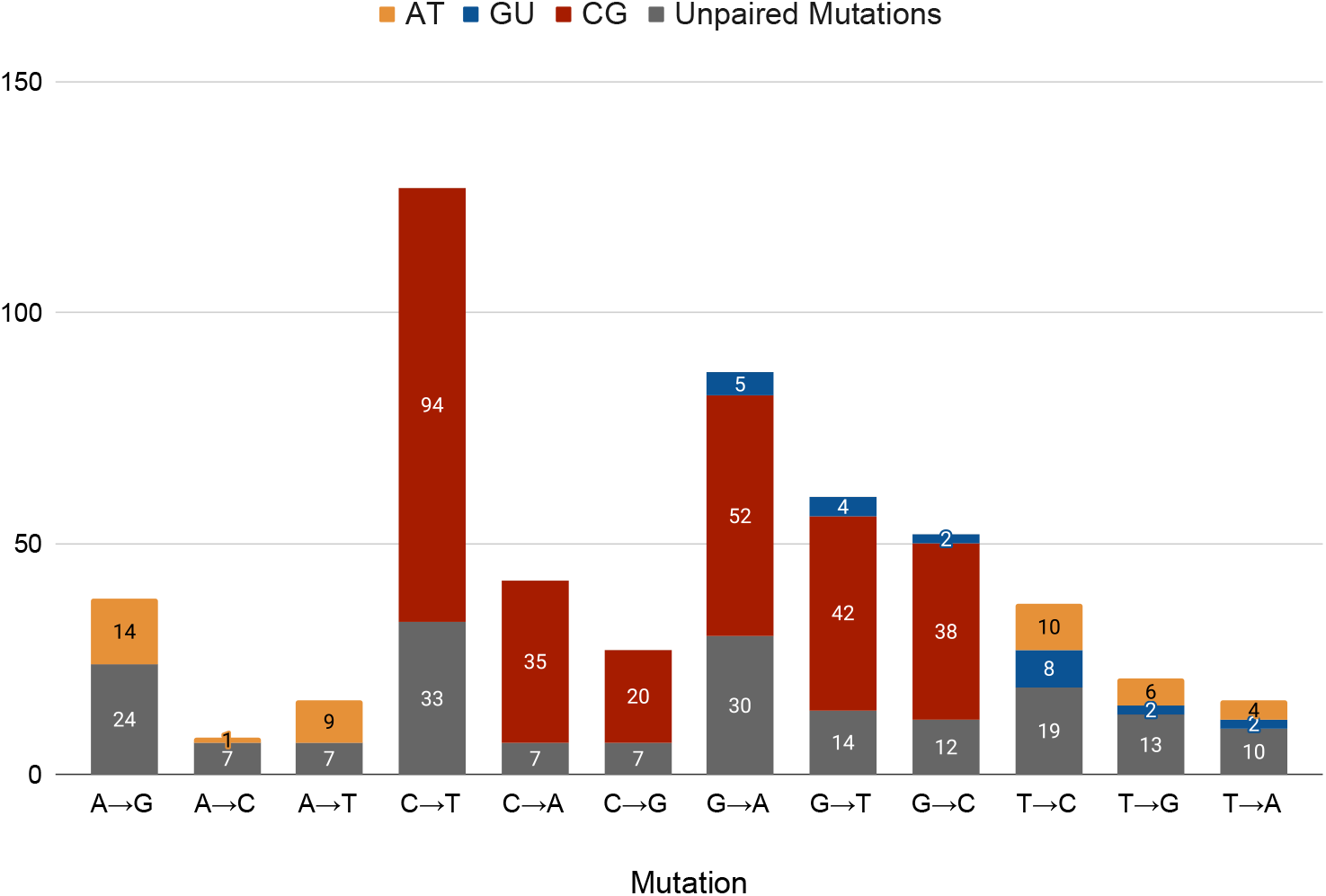
Number of observed substitutions, categorized by pairedness with pairings further subdivided by type. Substitutions are ordered first by the nucleotide that was substituted and then the resulting substitution. Bar segments are colored to show the base pair in the secondary structure when the substituted nucleotide was paired, and gray when the mutated nucleotide was unpaired.

Similarly, we do not see enhanced conversion of GU pairings into GC/AU ones, as might also be expected. There are 974 wobble pairings in *S* (8.7% of 11,208) but GU is only 6.6% of the 348 pairings mutated to form *S*^*′*^. This is surprising as both transitions preserve pairing potential, and moreover would yield a base pair which is more thermodynamically stable.

Hence, we see a bias towards mutations in paired positions, particularly GC pairings, rather than away.

Of the Watson-Crick pairings that are mutated, only 1/3 could form a wobble pairing. And there is no enhanced conversion of wobble pairings under mutation.

Thus, the observed substitutions follow the mutational pattern of TAM and remarkably, these are not selectively removed to preserve structural integrity. This in in line with the empirical estimates of fitness effects by Li et al. (2016), which showed that for the single-copy of arginine-CCU tRNA in yeast, overall fitness decreased more at unpaired nucleotides than at paired ones.

### 4.3 Comparison with neutral neighbors

When considering the distribution of alleles encoded in *S*^*′*^, we found that mutations arising from TAM are likely to substantially compromise tRNA fitness by destabilizing molecule structure. In particular, 42.7% of observed one-point mutations disrupt pairings. Yet, it is well-established that secondary structure conservation is a critical factor in tRNA macroevolution — so much so that MFE-preservation is considered a reliable proxy for fitness (Fontana and Schuster, 1998; Reidys et al., 2001; Aguirre et al., 2011). Here we show that even when our dataset is restricted to the neutral neighbors, clear deviations from macroevolution expectations are seen.

Neutrality under thermodynamic optimization is a strong constraint. Although 57.3% of substitutions from *S* to *S*^*′*^ preserve pairing potential, only 37.3% are MFE-preserving. Interestingly, nearly all the difference is from the unpaired positions. The net result is that mutations still occur in paired nucleotides slightly more often than unpaired: 57.1% versus 42.9%. This is less than the substituted nucleotides for the full dataset (65.5% paired) but considerably more than those for the related neutral neighbors (32.6% paired). Thus, under this setting, all these observations agree with those in the Section *“Assessing preservation of pairing potential*.*”*

Moreover, under this scenario, we still find that GC pairings are mutated more than expected from the (initial) nucleotide distribution of MFE-preserving genes. This is again consistent with our TAM-bias model, but not with thermodynamic stability. In contrast, the initial distribution, which closely resembles the full one, favors GC over AU and GU pairings roughly 8:3:1. Hence, we see a bias towards the thermodynamic stability (with GC being by far the most stable pair, followed by AU) expected by macroevolution (Forster et al., 2006; Waldispühl et al., 2008) in the initial distribution, which is then being diminished by TAM-biased microevolution.

To quantify this bias further, recall that *M* denotes the MFE-preserving alleles from *S*^*′*^ and *α*(*m*) the corresponding gene in *S* for each *m* ∈ *M*. The set of all related neutral neighbors for *m* is 𝒩_*m*_, and 𝒩 = ⋃_*m*∈*M*_ *N*_*m*_. For each *m* ∈ *M*, we now consider the relative thermodyanmic stability *f*_*m*_ for *m* and *α*(*m*) in comparison to *N*_*m*_. As described in *Materials and Methods*, the *f*_*m*_ value ranges from 0 to 1 with higher being less favorable.

As seen in Figure 7 in *Appendix*, the values for the genes *α*(*m*) are fairly narrowly clustered with a mean of 0.415 and standard deviation of 0.084. In contrast, those for the alleles *m* are distributed over the full range with a (mean, std dev.) = (0.563, 0.258).

Hence, although GC pairs are clearly favored in the consensus genes, they are not the most thermodynamically stable in their neutral neighborhood. Interestingly, this may be indicative of mutational robustness (Gabzi et al., 2022) and/or evolvability (Wagner, 2023). In contrast, the observed neutral neighbor alleles tend to be among the less energetically favorable possibilities, once again demonstrating a departure from the macroevolutionary expectation for favoring stability. Thus, even under this context, observed mutations are not preferentially biased toward maintaining structural stability; instead, structural stability is negatively affected.

Finally, and to further reiterate such claim, we consider the positional distributions for *M* and *N* across the different tRNA substructures. As illustrated in Figure 4, the latter closely resembles findings for neutral neighbors from macroevolution studies (Fontana and Schuster, 1998; Reidys et al., 2001). However, there are key differences for the former microevolution distribution of substituted nucleotides.

**Figure 4.**
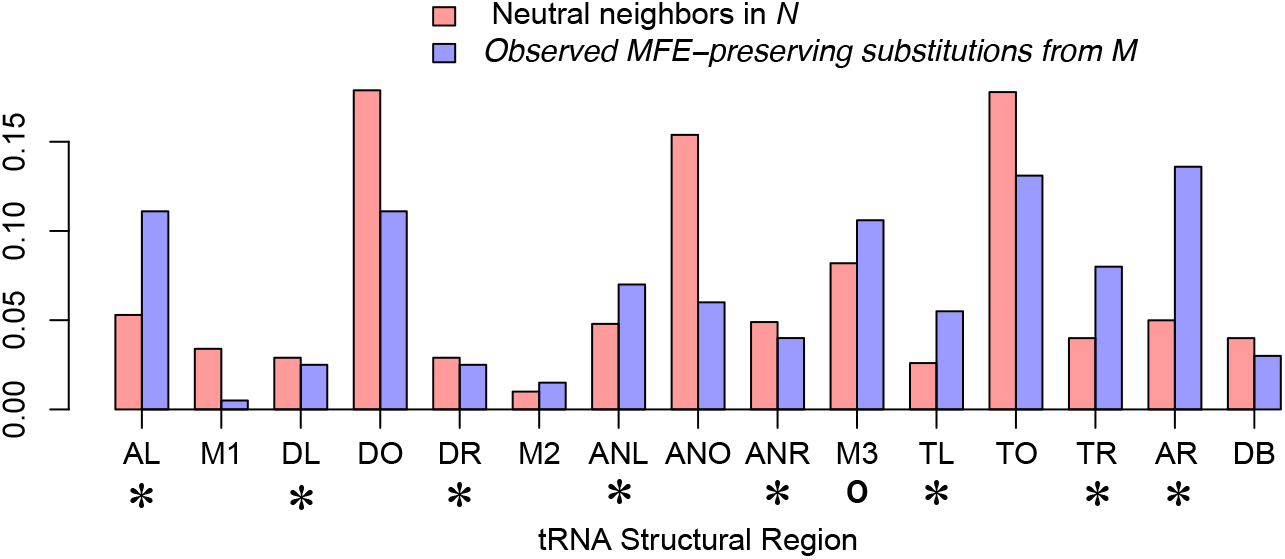
Distribution of observed MFE-preserving substitutions from *M* (blue) compared to all related neutral neighbors in 𝒩 (red). Components of the tRNA secondary structure are labeled as in Figure 1 with paired regions marked by *. Note the enhancement of paired nucleotides that were substituted in *M*, particularly the acceptor stem, while mutations in hairpin loops, especially the anticodon one, are depressed. Conclusions about the variable arm (M3) are confounded since it can contain both kinds of sites depending on the specific tRNA.

Observed MFE-preserving mutations occur more frequently in paired nucleotides, most notably in the acceptor stem (AR/AL) but also the 3’ side of the T-arm (TR). The variable arm portion of the multibranch loop (M3) is also enhanced, but conclusions are confounded since it can contain paired as well as unpaired sites depending on the particular tRNA. In contrast, substitutions in all three hairpin loops (DO, ANO, TO) are depressed, most notably for the anticodon.

As a common identity element shared among most tRNAs, the three anticodon nucleotides are an essential component of a functional tRNA molecule, so it is hardly surprising if they are less mutable than might otherwise be expected under thermodynamic predictions. However, anticodons are also known to evolve rapidly to meet new demands (Yona et al., 2013), and hence were included in our TAM-biased model. Although the anticodon itself represents *<* 1*/*2 of its hairpin loop, it was not *a priori* clear that there would be a discernible difference at this level of structural resolution.

### 4.4 Agreement with experimental fitness assays

We have demonstrated that the distribution of alleles in *C. elegans* tRNA is described by our new TAM- biased Markov substitution model but deviates from the expectation that disruptions to structure are the least tolerated mutations, even when restricted to neutral neighbors. We now show that this distribution, and by extension our conclusions, accords with fitness tRNA studies (Yona et al., 2013; Li et al., 2016; Domingo et al., 2018).

Recall that their experiments use the single-copy arginine-CCU gene from *S. cerevisiae*, so are insulated from the confounding effects when multiple copies of the same isoacceptor are present. Nonetheless, as seen in Figure 5 and detailed below, we find good agreement between the site-fitness distribution for all one-point mutants obtained by Li et al. (2016) and the location of substitutions in *C. elegans* alleles.

**Figure 5.**
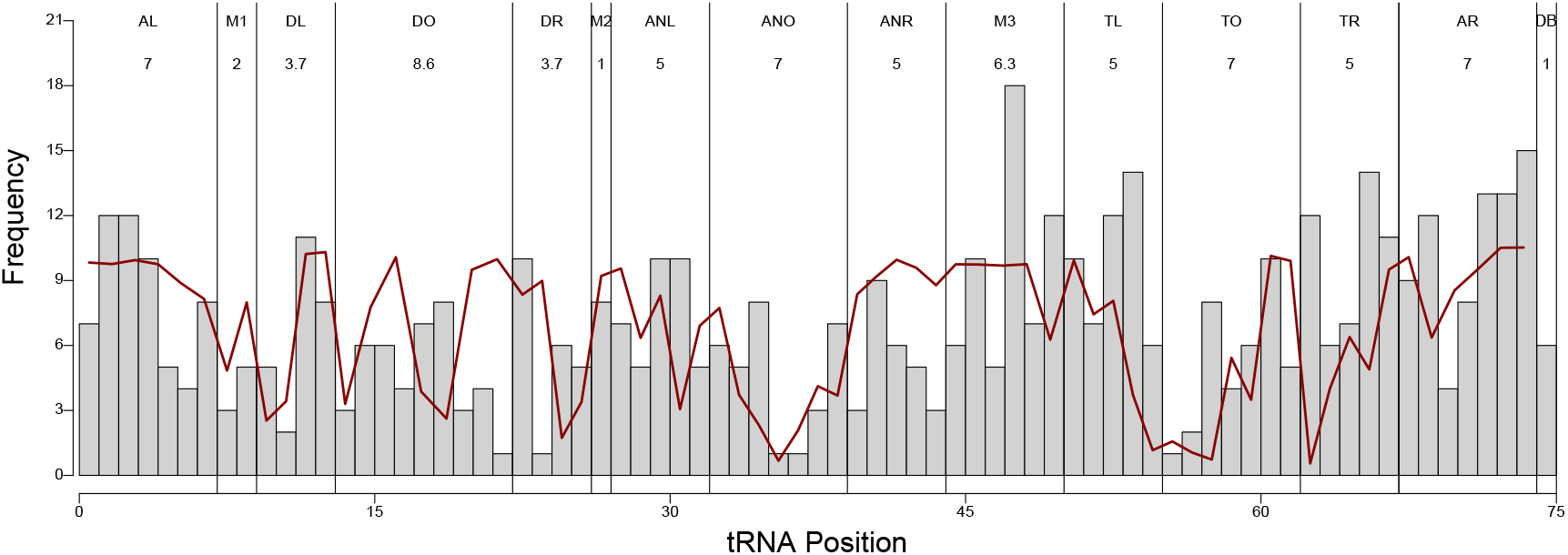
Comparison of substitution frequency over 531 *C. elegans* tRNA alleles (represented with gray bins) with mean site-fitness (Li et al., 2016) across all one-point mutants for the *S. cerevisiae* arginine-CCU tRNA (represented with the red line segments). Substructure components are labeled as in Figure 1 along with their average length, which was used for the site normalization.

To begin, we see that all *C. elegans* sites are mutated, but that lower frequency ones are often at (or near) a *S. cerevisiae* fitness minimum. Specifically, we observed a trend correlation of ∼0.5 when comparing the two datasets using a 3-nucleotide rolling average, and a point-wise correlation of around ∼0.4. We note that, for many reasons, a perfect correlation is not expected as, for example, this analysis includes different tRNA genes which have distinct identity elements, and thus different constraints.

In particular, the lowest frequency/fitness sites are seen in hairpin loops, specifically the anticodon and the 5’ end of the T-loop. The former agrees with both selection to preserve the anticodon as well as experimental evidence that it can evolve rapidly when needed (Yona et al., 2013), while the latter is conserved as part of the internal promoter B-box region (Li et al., 2016). There are also proximal frequency/fitness lows in the D-stem which are not (yet) explained by functional conservation. However, this conservation may be driven by the D-stem’s involvement in the A-box internal promoter essential for tRNA transcription (Mitra et al., 2015).

Conversely, regions with higher average frequency, i.e. both sides of the acceptor stem as well as the variable portion of the multiloop, also have high average fitness. This also echoes results from Domingo et al. (2018), whose dataset is built from 14 existing mutations of which 13 are from these regions.

Broadly, then, these two very different datasets agree; the acceptor stem and certain other paired sites are more mutable than expected under the (thermodynamic) neutral neighbor hypothesis, while the hairpin loops are much less so. These findings — aligned with our TAM-biased model — underscore the surprising robustness of many paired positions to mutations while also highlighting the influence of biochemical and other constraints beyond the secondary structure on the loop composition.

## 5 Conclusions

It is well-established that tRNAs are subject to strong purifying selection that maintains their structural integrity over macroevolutionary time scales. This is most apparent in the conservation of secondary structure, as evidenced by co-varying base pairs. Purifying selection is also evident at the microevolution time scale, as tRNA gene bodies exhibit substantially reduced rates of substitution compared to their flanking regions.

However, our results reveal that while the distribution of alleles in extant *C. elegans* strains is consistent with our model of TAM bias, it is not consistent with the expectations of thermodynamic neutrality for secondary structures. This shows that tRNAs in natural populations carry substantial levels of structure-destabilizing mutations, which may be tolerated but nevertheless likely induce meaningful fitness costs. This finding supports the conclusions in Thornlow et al. (2020), which claims that eukaryotic genomes carry substantial mutation load at tRNA genes.

This highlights a disconnect between scales. At the microevolutionary level, tolerable mutations often disrupt secondary structure, as demonstrated by empirical observations of fitness effects in yeast experiments. Yet, at the macroevolutionary level, selection pressure preserves canonical pairings. The resolution of this micro/macro dichotomy is likely to involve a variety of factors including the role of compensatory mutations (Kern and Kondrashov, 2004; Meer et al., 2010) and the interaction of isoacceptors (Yona et al., 2013) as well as the multi-copy nature of most tRNA genes (Bloom-Ackermann et al., 2014).

Beyond this, our new population model may be applicable in other contexts. The key assumptions are high gene conservation and short evolutionary time to the most recent common ancestor. These enable the use of a consensus sequence as the ancestral approximation and the encoding of the allelic distribution in a single compound mutational variation ‘descendant’.

It may also be possible to integrate other mutational biases into a Markov model through careful rate matrix parameterization. This naturally leads to questions of model identifiability, whose resolution may reveal new biological insights.

## 6 Materials and Methods

In this work we explore how TAM’s mutational patterns affect the the secondary structure of tRNA molecules by analyzing tRNAs from a *C. elegans* population. To do this, we introduce a continuous Markov model that describes the distribution of alleles under mutational bias across a population. We then apply this new model to analyze the distribution of tRNA genes across extant *C. elegans* strains.

Typically (Yang, 2006; Felsenstein, 2003), Markov models are used for phylogenetic inference across species. This is effective since the populations have diverged enough that it is possible to reconstruct their evolutionary relationships in a gene tree. In contrast, we seek not to characterize the relationship among the *C. elegans* alleles but only to describe the distribution of mutations. In doing so, we show that they are consistent with our microevolution TAM-bias substitution model but not with macroevolution expectations of thermodynamic neutrality for secondary structure conservation.

### 6.1 Model development

Our model formulation is possible under a strong purifying selection and high mutation rate over a short evolutionary timescale — exactly the situation for tRNA microevolution. As we show below, these conditions allow us to capture nearly all of the observed allelic variation via a continuous Markov substitution process.

#### 6.1.1 Modeling allele formation within a population

Let *N* be a population of *n* individuals and, for each individual *i* ∈ {1, 2, …, *n*}, let *S*_*i*_ be the concatenated collection of *k* genes under similar selection pressure, e.g. all tRNAs in extant *C. elegans* strains. Let *S* be the corresponding sequence for the most recent common ancestor of *N*, and denote the consensus sequence for the *S*_*i*_ as *S*.

We assume that the *k* genes are highly conserved and the evolutionary time from *S* is relatively short. Thus, the sequences *S*_*i*_ are all close to each other, and also to *S*. In this scenario, although not necessarily others (Czech et al., 2018; Trudeau et al., 2016), it is reasonable to approximate *S* by *S*.

To formulate our microevolution population model, we impose the following two requirements. Moreover, we show in *Results* that, for the tRNA microevolution situation being considered, these conditions are actually fairly mild.

1. An allele differs from *S* by one nucleotide substitution.
2. Two alleles in distinct individuals cannot differ in the same site.

Alleles meeting (i) are, by definition, one-point mutants, and are often the individual steps in macroevolution models (Saks et al., 1998). We refer to those satisfying (ii) as *distinct-site* mutants. Under these two conditions, we can encode all mutational variation from the compliant alleles in a single compound sequence.

Specifically, we introduce the *mutational variation sequence S*^*′*^ which is constructed by incorporating all observed substitutions in alleles meeting conditions (i) and (ii) into the consensus sequence *S*. As illustrated in Figure 6, this guarantees a one-to-one correspondence between sites where *S*^*′*^ differs from *S* and all distinct-site, one-point mutations observed in the population.

**Figure 6.**
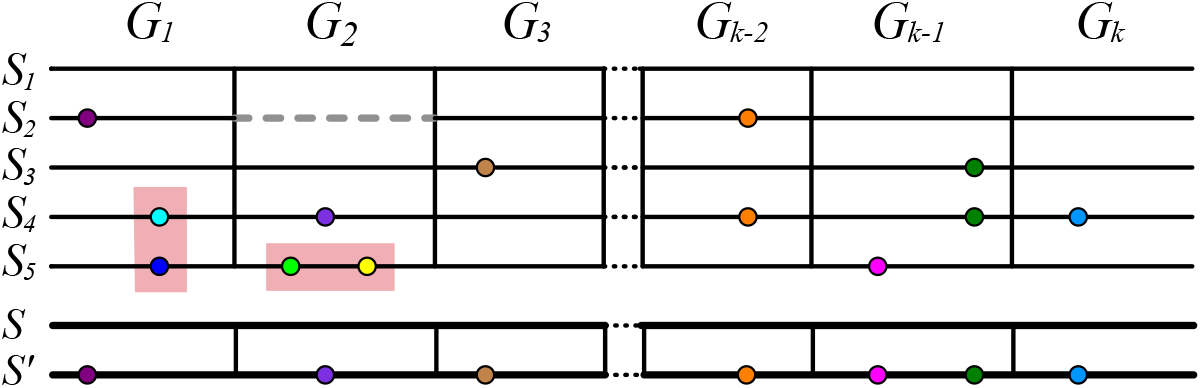
Schematic of mutational variation sequence construction. From top to bottom: Each line *S*_*i*_ represents the *k* concatenated genes *G*_*j*_ for an individual *i* in the population *N* = {1, …, 5}. Every gene is not necessarily present in each individual; dashed gray lines denote a missing gene. Colored dots indicate nucleotide substitutions with respect to the consensus sequence *S* (shown below *S*_5_). Highlighted in red, we see that *G*_2_ in *S*_5_ violates condition (i) and *G*_1_ violates (ii) in *S*_4_ and *S*_5_. By treating alleles violating these two conditions as missing genes, the remaining allelic variation can be encoded in the new compound sequence *S*^*′*^ by substituting into *S* each distinct-site one-point mutation.

**Figure 7.**
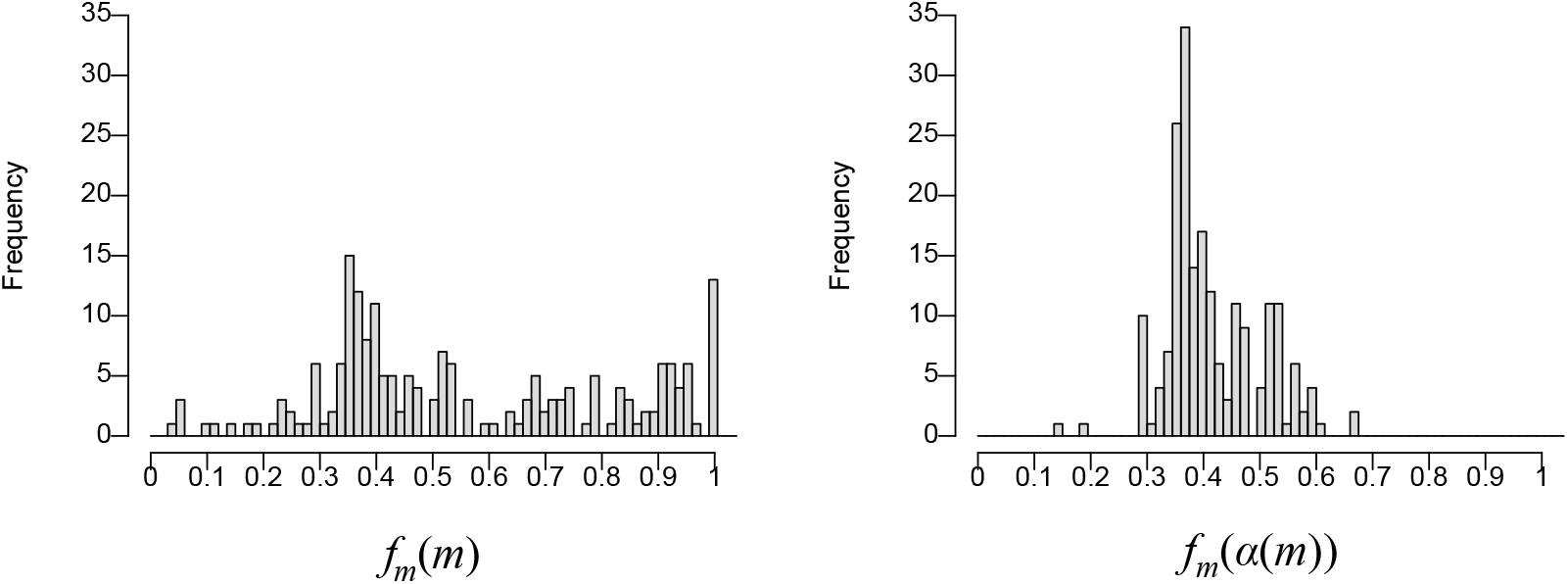
Relative thermodynamic stability *f*_*m*_ for the 198 observed neutral neighbors (left) and corresponding tRNA gene (right). Approximately half of the observed substitutions in the *C. elegans* population are disproportionately less stable than other related neutral neighbors, but no such skew is seen for the consensus tRNA genes approximating the ancestral sequence.

By integrating all those mutations into a single sequence, we can model allele formation without requiring a phylogenetic tree. This is particularly useful as inferring the true evolutionary relationship between individuals within a population can be very challenging (De Maio et al., 2015). Instead, we describe it simply through a nucleotide substitution process from *S* to *S*^*′*^.

Moreover, since multiple mutations within the same gene occur in different alleles, we strengthen the site independence for the Markov model. Hence, we parameterize our model with a 4 *×* 4 rate matrix for nucleotide substitutions, rather than the more complicated dinucleotide ones (Allen and Whelan, 2014) which model RNA pairing covariation for macroevolution analyses.

#### 6.1.2 Modeling the distribution of substituted nucleotides under TAM

As described in the *Background*, TAM’s characteristic signature results in an asymmetric increase in *C* → *T* and *G* → *A* mutations on the coding strand. We hypothesize that the exchangeability asymmetry caused by TAM results in a bias of the distribution of substituted nucleotides, which we call the *TAM bias*.

To formulate this mathematically, let *π* = (*π*_*A*_, *π*_*C*_, *π*_*G*_, *π*_*T*_) be the nucleotide distribution of the consensus sequence *S* (an approximation of *S*, the ancestral sequence). With no mutational bias, one would expect that the distribution of substituted nucleotides follow *π*.

To adjust for TAM, we introduce the *TAM bias vector τ* = (*τ*_*R*_, *τ*_*Y*_) ∈ [0, 1]^2^. We hypothesize that the nucleotides in *S* that mutated to form *S*^*′*^ follow the *TAM bias distribution* given by:

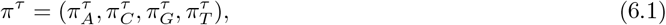

where

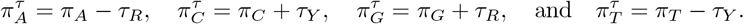

In other words, the TAM bias vector *τ* adjusts the distribution of substituted nucleotides to account for differences in transition rates within purines (*R*) and within pyrimidines (*Y*). Hence, 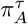 is the proportion of *A*’s in *S* that mutated to another nucleotide to form *S*^*′*^ relative to all observed distinct-site, one-point substitutions.

#### 6.1.3 Modeling a TAM-biased substitution process

Our proposed TAM-biased distribution describes how the distribution of substituted nucleotides is altered from the root nucleotide distribution in *S*. However, to model allele formation within a population under TAM, we need to account for the mutation process itself. For this, we present a continuous Markov substitution model that incorporates the TAM bias to describe the production of *S*^*′*^ from *S*.

Recall that for a Markov process over an edge of length *t*, the joint distribution of nucleotides in the ancestral and descendant sequences is given by *P* = diag(*π*) exp(*Qt*). In our context, the entry *p*_*ij*_ of *P* represents the probability of a site in *S*^*′*^ differing from *S* by a substitution from *i* to *j*. To parameterize this new model *P*, we must propose a TAM-biased instantaneous rate matrix *Q*. Recall also that, typically, one entry of the matrix is set to 1 to avoid scaling redundancy with the edge length. We note that the rate matrices presented here are not-normalized form for simplicity, but all computations are done under normalization as described in the *Background*.

We begin with a strand-symmetric model (*sym*) which assumes that a substitution is indistinguishable from its Watson-Crick complement. This accounts for the background noise, and its matrix has 5 free parameters with:

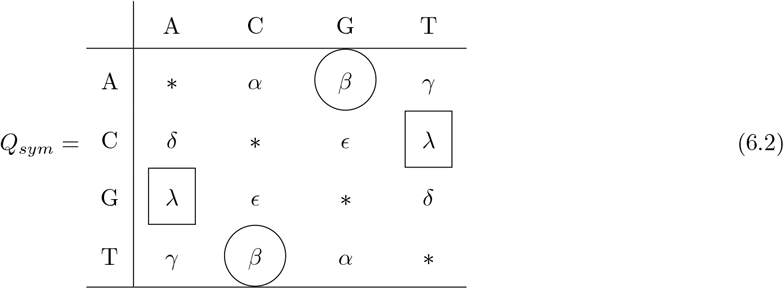

where *α, β, γ, ϵ, λ* ≥ 0 and *δ* = 1. Note that by accounting for the edge length, the model has 6 free parameters.

However, a characteristic of TAM is the asymmetry in transitions observed in the coding strand. Hence, we break the strand symmetry between substitutions *A* → *G* with *T* → *C* (circled in *Q*_*sym*_), and *G* → *A* with *C* → *T* (squared in *Q*_*sym*_), yielding a transition-asymmetric rate matrix

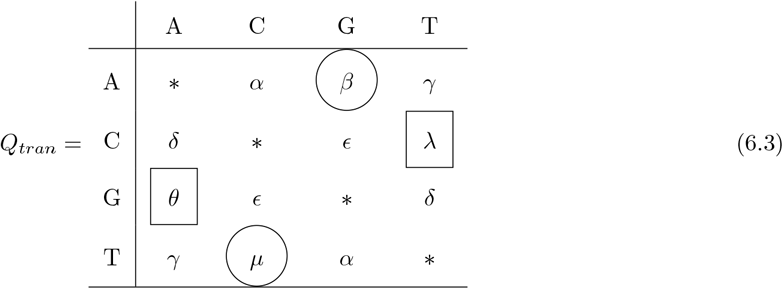

with 7 free parameters. Hence, *Q*_*sym*_ is a special case of *Q*_*tran*_ with *θ* = *λ* and *µ* = *β*, and this model has 8 free parameters when accounting for the edge length.

Next, we incorporate the TAM bias model for the distribution of substituted nucleotides by adding restrictions to the *Q*_*tran*_ rate matrix entries. More specifically, we require that

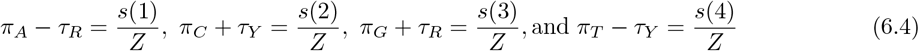

where *s*(*i*) = ∑ _*j*≠*i*_ *p*_*ij*_ is the sum of the off-diagonal entries in each row of *P* with *Z* = ∑ _*i*≠*j*_ *p*_*ij*_ as the normalizing factor. Thus, the left-hand sides come from Equation (6.1), whereas the right ones follow from the continuous Markov model.

We remark that these equations do not increase the number of model parameters. The entries of *π* are needed to compute *P* for every rate matrix, and hence do not count against any particular model. Additionally, the TAM bias vector *τ* = (*τ*_*R*_, *τ*_*Y*_) is estimated from the flanking regions, which are subject to TAM but not strong selection pressure. Hence, since *τ*_*R*_ and *τ*_*Y*_ are independent from the tRNA alleles, they are not counted as model parameters either.

As a consequence, we improve the model parameterization under the constraints in Equations (6.4) by identifying linear dependencies among the *Q*_*tran*_ rate matrix entries using a computer algebra package. More precisely, given the high conservation of tRNAs, we expect a very slow substitution process, implying that the entries of *Qt* are quite small. Thus, we approximate exp(*Qt*) ≈ *I* + *Qt*, where *I* is the 4 *×* 4 identity matrix. Under this approximation, we represent Equations (6.4) as a homogeneous linear system which can be solved using Macaulay2 (Grayson and Stillman, 2009).

As a result, we obtain the linear dependencies given below for the transversions *δ* and *γ* as well as the transition *θ*. We then define *Q*_*TAM*_ to be the rate matrix obtained from *Q*_*tran*_ using these relationships. This yields a total of 4 free parameters for the new TAM-biased rate matrix: the 3 transitions *β, λ*, and *µ* along with the 2 transversions *α* and *ϵ* – yielding 5 free parameters for the model.

We note that the choice of dependent variables among the matrix parameters was determined by Macaulay2 (version 1.25.06); any alternate formulation of the relationships would yield an equivalent model parameterization although a different form for the *Q*_*TAM*_ rate matrix. The linear dependencies used here (which were manually derived from Macaulay2’s initial output shown in the Appendix) are:

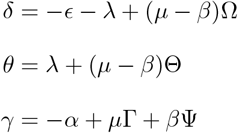

where

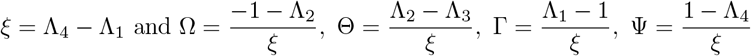

for

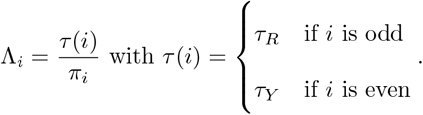

Observe that the coefficients Ω, Θ, Γ, and Ψ are real numbers defined solely by the TAM bias vector *τ* and the root nucleotide distribution *π*. As a function of *π* and *τ*, they are unbounded and defined almost everywhere (except for a set of measure zero, that is, a negligible subset of the parameter space).

We remark that Ω ≠ 0. Hence, the inequality *µ* ≠ *β* must hold; otherwise *δ <* 0, which violates the definition. Likewise Θ ≠ 0 for almost all *π* and *τ* . But since we have already concluded that *µ* ≠ *β* when Ω is defined, then generically *θ* ≠ *λ* as well. This shows that satisfying Equations (6.4) requires breaking the transition strand symmetries.

We show in *Results* that among the five rate matrices considered — strand-symmetric (*sym*), transition-asymmetric (*tran*), and TAM biased (*TAM*) along with general Markov (*GM*) and general time-reversible (*GTR*) — the new TAM-biased Markov substitution model best fits the allelic distribution in our *C. elegans* tRNA dataset.

### 6.2 Data analysis

Our dataset was generated from two publicly available databases: the Genomic tRNA Database (GtR-NAdb, v2.0) (Chan and Lowe, 2008, 2015) and the *C. elegans* Natural Diversity Resource (CeNDR, release 20180527) (Cook et al., 2016; Crombie et al., 2023). The starting point was the reference genome (release WBcel235/ce11) (Harris et al., 2019) for the universal laboratory strain N2.

GtRNAdb identifies 581 functional tRNAs in the N2 nuclear genome, and CeNDR provided variant data for 330 *C. elegans* wild isolates collected from around the globe. We then retrieved 677 single nucleotide variants (SNVs) for the tRNA genes from the hard-filter variant call format (vcf) file using custom Python scripts.

Secondary structures for all tRNA sequences were obtained from `tRNAscan-SE` (version 2.0.11 (Chan and Lowe, 2019)) using the default parameters. As illustrated in Figure 1, nucleotides were classified as “paired” or “unpaired” according to these structures. We note that, although pictured, the Tail was not included in the analysis.

There were 16 SNVs which were found by `tRNAscan-SE` to occur in an intron; these were excluded from further analysis. Hence our *C. elegans* microevolution dataset consists of 661 variants for 581 tRNA genes across a population of 331 strains.

We also collected data for the 40 nucleotides upstream and 40 downstream of each SNV and N2 gene. Since these tRNA flanking regions are unwound during transcription, they are subject to TAM (Thornlow et al., 2018). However, they are not under strong selection pressure, making them ideal for estimating the TAM bias vector *τ* = (*τ*_*R*_, *τ*_*Y*_). Under our TAM bias hypothesis, subtracting the of observed distribution substituted nucleotides from the initial nucleotide distribution should yield *π* − *π*^*τ*^ = (*τ*_*R*_, −*τ*_*Y*_, −*τ*_*R*_, *τ*_*Y*_). As reported in *Results*, the positive and negative values are in good agreement, yielding a robust estimate for *τ* .

#### 6.2.1 Testing TAM-biased distributions

Our model for TAM bias from Equation (6.1) proposes that the substituted nucleotides are distributed as *π*^*τ*^ where 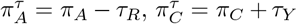, etc. To validate this hypothesis for the TAM-biased rate matrix, we compute a *p*-value using the `xmonte` function from the XNomial R package (Engels, 2015). This function assesses whether the observed substitutions conform to the estimated multinomial distribution. Specifically, it examines 3,000 random outcomes to estimate the probability that a sample deviates from the expected distribution by at least as much as the observed data.

We later apply the same approach to test for the preservation of pairing potential. To do this, we classify the nucleotides of each tRNA gene as either paired (p) or unpaired (u) according to the `tRNAscan-SE` secondary structure. This separates *π*, the nucleotide distribution for *S*, by pairedness into

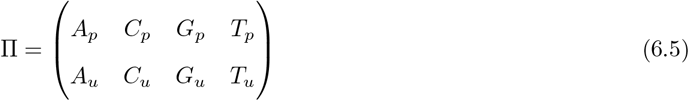

where *X*_*p*_ and *X*_*u*_ denote the proportion of paired and unpaired nucleotides respectively for *X* ∈ {*A, C, G, T* }. Marginalizing the rows of Π reconstitutes *π*, while column marginalization yields the pairedness distribution, denoted Π_*pu*_ = (Π_*p*_, Π_*u*_).

We also consider the distribution of base pairs, denoted *ρ* = (*ρ*_*GC*_, *ρ*_*AT*_, *ρ*_*GT*_), where *ρ*_*GT*_ is the proportion of GU pairings in the tRNA secondary structures. It follows that

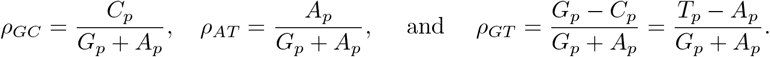

Under our TAM bias hypothesis, the distribution of substituted nucleotides by pairedness would be

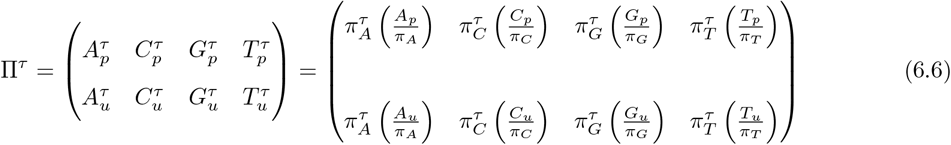

where, for instance, 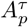 is the proportion of one-point mutants in *S*^*′*^ that differ from *S* by a substitution which changed a paired *A*. Row and column marginalization of Π^*τ*^ yield *π*^*τ*^ and 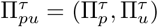 respectively.

Analogously, 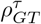 represents the proportion of substitutions that changed either the *G* or the *T* in what would have been a wobble *GU* pairing. Consequently, the components of *ρ*^*τ*^ are

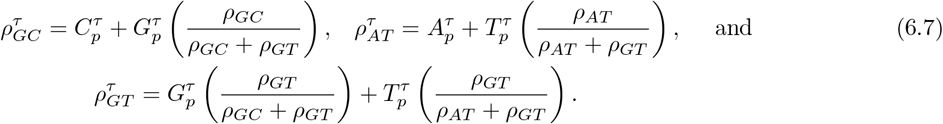

#### 6.2.2 Assessing Markov substitution rate matrices

To compare the five possible exchangeability matrices considered, we computed for each the three most common information criteria for model selection (Kalyaanamoorthy et al., 2017): the Akaike (AIC) (Akaike, 1974), the corrected Akaike (AICc) (Burnham and Anderson, 2002), and the Bayesian (BIC) (Schwarz, 1978). We found that the differences between the first two were always negligible due to the length of the consensus sequence relative to the number of mutations. Hence, we focus on the AIC and BIC.

Loosely speaking, the lower the estimate, the ‘better’ the model. Each criterion depends on the log-likelihood (LH) of the model being ‘penalized’ by the number of parameters. The LH function is given by

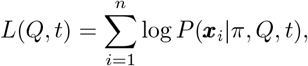

where *π* is the nucleotide distribution of *S, n* is the sequence length of *S*, ***x***_*i*_ is the site pattern of *S* and *S*^*′*^ at site *i, Q* is the rate matrix, *t* are the expected number of substitutions between *S* and *S*^*′*^, and *P*(***x***_*i*_|, *π, Q, t*) is the conditional probability of ***x***_*i*_ given *π, Q*, and *t*.

To compute the LH function and estimate the maximum likelihood estimator (MLE) for all parameters, we again use the approximation exp(*Qt*) ≈ *I*+*Qt*. We then used MATLAB’s (version 25.2, R2025b (MathWorks, 2024)) `fmincon` function to estimate *Q* and *t* MLEs using 1000 different random starting points.

We also computed 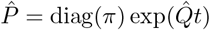 using the MLE estimates for each parameterization and compared the results to the data. The corresponding *p*-values for such comparisons were then obtained using the `xmonte` function.

#### 6.2.3 Analyzing thermodynamic neutrality

We also compared observed distributions against a background of neutral neighbors. Recall that these are alleles which are both one-point mutants and MFE-preserving. For consistency, we consider only observed alleles which are also neutral neighbors. The necessary thermodynamic predictions, i.e. the MFE score and its associated secondary structure, are computed with `RNAfold` (version 2.6.2 (Lorenz et al., 2011)) under default parameters.

More precisely, let *P* be the set of all distinct-site, one-point mutant alleles used to generate *S*^*′*^, and *R* the set of tRNA genes in *S*. For *p* ∈ *P*, let *α*(*p*) be its corresponding *r* ∈ *R*. Then *p* ∈ *P* is MFE-preserving, and hence is a neutral neighbor, when it has the same `RNAfold` predicted secondary structure as *α*(*p*).

Let *M* ⊆ *P* be MFE-preserving, i.e. all observed neutral neighbors. For *m* ∈ *M*, let *N*_*m*_ be the set of all neutral neighbors of *α*(*m*), and 𝒩 = ⋃ _*m*∈*M*_ *N*_*m*_. Since *m* ∈ *N*_*m*_, *M* is a subset of *N* . We determine *M* and 𝒩 computationally, and consider how they differ.

First we consider whether the substituted nucleotides were paired or not, according to the classification from the `tRNAscan-SE` secondary structure for *α*(*m*). Next we consider the thermodynamic stability of *m* and *α*(*m*) relative to *N*_*m*_ over all of *M*.

This is done by ranking the set *N*_*m*_ ∪ {*α*(*m*)} by increasing MFE score with the lowest ranked as 1. The ranking is dense (García-Lapresta and Martínez-Panero, 2024); ties share the same rank, and the next rank is assigned the next integer. For instance, if three sequences tie for the 5th lowest MFE score, all are assigned rank 5 with the next one(s) ranked 6. Rankings are then normalized by dividing by the total number of ranks for *α*(*m*). This defines a function *f*_*m*_ : *N*_*m*_ ∪ {*α*(*m*)} → (0, 1] for each *m* ∈ *M* where values near 0 are the most thermodynamically stable while those near 1 are the least so. We will compare *f*_*m*_(*m*) and *f*_*m*_(*α*(*m*)) over all of *M*.

Finally, we consider the positional distribution of substitutions in *M* and 𝒩 . The tRNA structural regions are defined in Figure 1 and determined for *α*(*m*) based on the `tRNAscan-SE` secondary structure.

#### 6.2.4 Comparing with experimental fitness

We consider the mutability of each site across the 531 alleles in our *C. elegans* dataset. Since not all tRNAs have the same length, the sites in each allele were normalized to the average length of the corresponding substructure component from the 15 (excluding the Tail) listed in the table in Figure 1. The number of mutations for each normalized site can then be presented in a frequency histogram.

We compared this distribution to the mean site-fitness scores obtained by Li et al. (2016) for the *S. cerevisiae* single-copy arginine-CCU tRNA. The sites in the 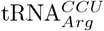 were also normalized as above before reporting the average of their 3 one-point mutant fitness scores. For display purposes, this mean value was linearly re-scaled from (0.685,1.017) to (1,10).

## 7 Data Availability

The data used in this work were obtained from and are available at the Genomic tRNA Database (GtR-NAdb, v2.0) (Chan and Lowe, 2008, 2015) and the *C. elegans* Natural Diversity Resource (CeNDR, release 20180527) (Cook et al., 2016; Crombie et al., 2023). The repository https://github.com/HectorBanos/TAM contains the concatenated gene tRNA sequence *S* (composed of all 531 tRNAs), and the population variation sequence *S*^*′*^, as well as code to reproduce the results in this paper.

## 8 Acknowledgments

This work was supported by the NSF-Simons Southeast Center for Mathematics and Biology (SCMB) through the grants National Science Foundation (NSF) DMS-1764406 and Simons Foundation/SFARI 594594. H.B. was also partially supported by NSF grant DMS-2331660. A.P. was also partially supported by NSF grant IOS-2319796. C.H. was also partially supported by NIH grant R01GM126554. The authors used a Large Language Model solely for spelling and grammar corrections; it was not used to generate original content.

## 9 Appendix

The output obtained from Macaulay2 by solving the linear system described in Section 6.1.3.

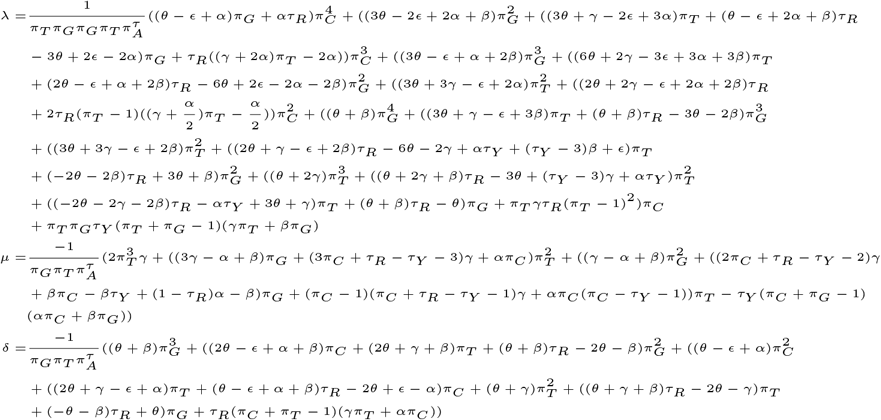

The dependencies of the *Q*_*TAM*_ matrix entries shown in Section 6.1.3 where derived manually from these.

## Footnotes

a Uracil (U) will be used when discussing RNA structures and thymine (T) for their DNA genes.

b Sequence pictured is *Bacillus subtilis* tRNA^Asp^ with structures adapted from Liu (2013).

c Although there are 20 standard amino acids and 64 distinct codons, most eukaryotic nuclear genomes encode hundreds of tRNAs.

d All data was analyzed at 10-digit precision. Due to rounding, reported values may not add up exactly.

